# Audio-visual concert performances synchronize an audience’s heart rates

**DOI:** 10.1101/2024.04.10.588486

**Authors:** Anna M. Czepiel, Lauren K. Fink, Mathias Scharinger, Christoph Seibert, Melanie Wald-Fuhrmann, Sonja A. Kotz

**Author notes:** Corresponding author Anna M. Czepiel Department of Psychology University of Toronto Mississauga 3359 Mississauga Road, Mississauga Canada.

## Abstract

People enjoy engaging with music. Live music concerts provide an excellent option to investigate real-world music experiences, and at the same time, use neurophysiological synchrony to assess dynamic engagement. In the current study, we assessed engagement in a live concert setting using synchrony of cardiorespiratory measures, comparing inter-subject, stimulus-response, correlation, and phase coherence. As engagement might be enhanced in a concert setting by *seeing* musicians perform, we presented audiences with audio-only (AO) and audio-visual (AV) piano performances. Only correlation measures were above chance level. In comparing time-averaged synchrony across conditions, AV performances evoked higher inter-subject correlation of heart rate (ISC-HR). However, self-reported engagement did not correspond to synchrony when averaged across music pieces. On the other hand, time-resolved analyses show that synchronized deceleration-acceleration heart rate (HR) patterns, typical of an ‘orienting response’ (an index of directed attention), occurred *within* music pieces at salient events of section boundaries. That is, seeing musicians perform heightened audience engagement at structurally important moments in Western classical music. Overall, we could show that multisensory information shapes dynamic engagement. By comparing different synchrony measures, we further highlight the advantages of timeseries analysis, specifically ISC-HR, as a robust measure of holistic musical listening experiences in naturalistic concert settings.

## 1. Introduction

People enjoy engaging in music, especially in live concerts^1–3^. While concerts provide an excellent option to explore music listening experiences, assessing this experience dynamically is methodologically challenging. One way to explore musical engagement over longer time scales is with synchrony of peripheral responses either across participants or to perceived musical structures. However, many different analysis techniques can be employed. Here, we compared different synchrony analyses to gauge their robustness. Using cardiorespiratory synchrony, the current study assessed engagement, defined as a listener’s real-time absorption in music^4–6^. As engagement might be enhanced in a concert setting by *seeing* musicians perform^7–10^, we presented participants with audio-visual and audio-only musical performances in a concert context.

Musical engagement can describe either involvement in musical activities (e.g., instrumental practice) or the time-varying mental state when listening to music^11^. Here, we refer to engagement as an absorbed state of heightened attention and affect, which can change on a second-by-second basis while listening to music^5,6,11–13^ (for a review, see^5^). Indeed, music is a particular driver of (regular) time-varying attentional focus due to its metrical structure/beat. Although musical engagement can be measured with continuous behavioural responses^12,13^, these ratings might distract from directly experiencing music. A promising alternative to measure engagement in a non-distracting way is to assess neural and peripheral physiological responses. One potential index is the orienting response: the engagement of attention, which is reflected in increased skin conductance and deceleration-acceleration patterns of respiration/heart rate^14^. Another method to objectively assess engagement is through physiological synchrony (e.g.,^2,15–18^), which assumes that increased engagement with music is more likely to evoke common time-locked responses across participants^4,19^. Synchrony can be between a participant’s response and a stimulus feature that represents how we process/respond to sound, like the auditory envelope (i.e., how sound intensity changes over time). More recently, spectral flux (spectrum changes over time) has been used as a more faithful representation of how humans respond to auditory stimuli^20–24^ as it represents both envelope and rhythmic information^24^ and is related to perceived higher activity (e.g., faster music) and fullness of sound^25^. Assessed by calculating the (Euclidean) distance between the spectrum of successive frames, spectral flux can also be assessed within specific frequency bands (sub-bands), which represent different rhythmic elements^26^. For example, in pop music, spectral flux of lower frequency bands can represent kick drum and bass guitar rhythms, while higher frequency bands can represent cymbals and hihat rhythms^27^. Synchronisation of participant responses to stimuli can be assessed using correlation or phase coherence, i.e., rhythmic alignment^4,24^. In this paper, we refer to these as stimulus-response correlation (SRC) and stimulus-response phase coherence (SRPC), respectively. A related synchrony approach is to assess similarity across multiple time-locked participant responses^28^. Similarly to stimulus-response synchrony, inter-subject synchrony can be calculated with correlation^17,29^ or using phase coherence; in this paper, we refer to these as inter-subject correlation (ISC) and inter-subject phase coherence (ISPC), respectively.

Growing evidence has related synchrony of neural responses to a participant’s engagement with stimuli. In correlational measures, SRC and ISC were related to engagement with movies^17,30,31^, speech ^32,33^, and music^2,4,13,19^. In phase coherence measures, higher neural synchrony occurred in engaging group discussions, indicating a potential marker for shared attention mechanisms^34^. Although phase synchrony in speech was attributed to stimulus intelligibility^35^, it was postulated that phase synchrony increases when engaging in stimuli. According to the Dynamic Attending Theory (DAT), internal oscillations adapt to external rhythms so that attending energy is optimised at expected time points^36,37^. Indeed, successful coordination in musical ensembles was related to phase synchrony in neural^38^ and heart^39^ rhythms (see also^40,41^). Recent work shows coupling of cerebral activity occurs when experiencing similar emotional response to live music concerts^15^. However, most synchrony research focused exclusively on neural responses, such as those measured via electroencephalography (EEG); more suitable approaches for a live concert audience could be to explore synchrony in peripheral physiological responses^2,16,42^.

Recent frameworks propose certain neural mechanisms – such as synchrony – might extend to cardiac, respiratory, gut, and pupil rhythms^43–45^. Promising results show cardiorespiratory ISC related to engagement with narratives and instructional videos^18,46^. In the phase domain, some research indicates that the alignments of breathing/heartbeats to external rhythms aid the processing of upcoming stimulus events^47,48^. In music, respiration has been shown to entrain to musical beats^49,50^, although heartbeats do not^51–53^. The current study aimed to further assess cardiorespiratory synchrony as an index of engagement in response to music performances in a concert audience.

Cardiorespiratory synchrony can be time-averaged across stimuli to assess engagement between conditions (e.g., attended/unattended conditions^18^). However, music listening is dynamic, where engagement, neural synchrony^12,13,54^, and physiological synchrony^16^ change over time. Previous research reported that engagement tends to peak at musical climaxes or structural boundaries^55^. Indeed, it seems beneficial for attentional resources to ‘latch onto’ structural boundaries: the brain naturally segments incoming information into more meaningful units to process them more efficiently^56,57^. According to the Generative Theory of Tonal Music (GTTM^58^), we ‘segment’ auditory events based on salient changes when several music features change simultaneously. The ability to segment music at such salient moments has seen consistent empirical support in behavioural ratings^59–62^. However, consciously rating engagement continuously distracts from a natural music listening experience. Recent research showed that salient musical section boundaries evoke neurophysiological synchrony, potentially reflecting heightened engagement^12,13,18^. Therefore, to complement the current investigation of engagement during audio-only (AO) and audio-visual (AV) music performance on a global level (time-averaged across pieces), we also tested time-varying engagement – indexed by physiological synchrony – between AO and AV music performances during salient moments, defined as section boundaries.

### The current study

Audiences might find concerts engaging due to not only listening to music but also seeing a musician perform^8,10^. To test this, concert audiences were presented with AO and AV performances of Western classical piano pieces. Musical pieces were selected as they represent a range of genres (Baroque, Classical-Romantic, 20^th^ century music), and characters, as well as range of tempi, harmonic and musical phrase structures. Presenting such a musical range was thought to evoke high engagement levels. Additionally, pieces were chosen as they may be considered typical pieces heard in live Western classical performances, though not known to the audience, thereby reducing familiarity effects. Based on Kaneshiro et al.^4^, dynamic engagement was assessed by calculating cardiorespiratory synchrony using both correlation and phase coherence. For correlation, we assessed heart and respiration rate, e.g., speed of breathing and heartbeats in terms of beats per minute (BPM). For phase coherence, we assessed the phase alignment of heartbeat cycles (i.e., whether heartbeats align in time) and breathing cycles (i.e., whether breathing aligns in time). Synchrony was then assessed both across (i.e., time-averaged synchrony) and within (i.e., time-resolved synchrony) musical pieces. First, we tested that time-locked responses were synchronized above chance level. Second, we hypothesised that time-averaged synchrony would be higher in AV than in AO musical performances. Third, we expected time-resolved synchrony to be higher at section boundaries (based on previous literature^12,13,16^). Further, we expect synchrony to be higher at section boundaries in AV than AO at salient moments in the music (interaction effect), based on literature showing that visual information enhances perceptual responses^63^ and that visual imagery heightens engagement^55^.

## 2. Method

Participants, stimuli, and procedures are identical to Czepiel et al.^7^ (see General Methods and Experiment 2). Key details of the procedure are outlined below. Data and code are available on Open Science Framework (OSF)^1^.

### Participants

The study was approved by the Ethics Council of the Max Planck Society and in accordance with the Declaration of Helsinki. Participants gave their written informed consent. Twenty-six participants attended the concerts. One participant was excluded due to missing physiological data, thus behavioural and physiological data of twenty-five participants were analysed. Participants included nine females, mean age of 51.08 years (SD = 15.48), who on average had 6.18 (SD = 8.20) years of music lessons and attended on average 13 concerts per year (M = 19.97, SD = 20).

### Stimuli

Participants were presented with AV and AO versions (see Figure 1a) of three Western classical piano pieces: Johann Sebastian Bach: Prelude and Fugue in D major (Book Two from the Well-Tempered Clavier, BWV 874), Ludwig van Beethoven: Sonata No. 7, Op. 10, No. 3, second movement (Largo e mesto), and Olivier Messiaen: *Regard de l’Esprit de joie* (No. 10 from *Vingt Regards sur l’enfant Jésus*). Both versions were performed by the same professional pianist, on the same piano, in the same concert hall. AO versions were recorded prior to the concert and were presented as recordings through high-quality loudspeakers, while AV versions were performed live by the pianist during the concert. A trained sound engineer checked that the sound level was equal across all stimuli; additional analyses of acoustic features showed these performances were comparable across concerts^7^.

**Figure 1.**
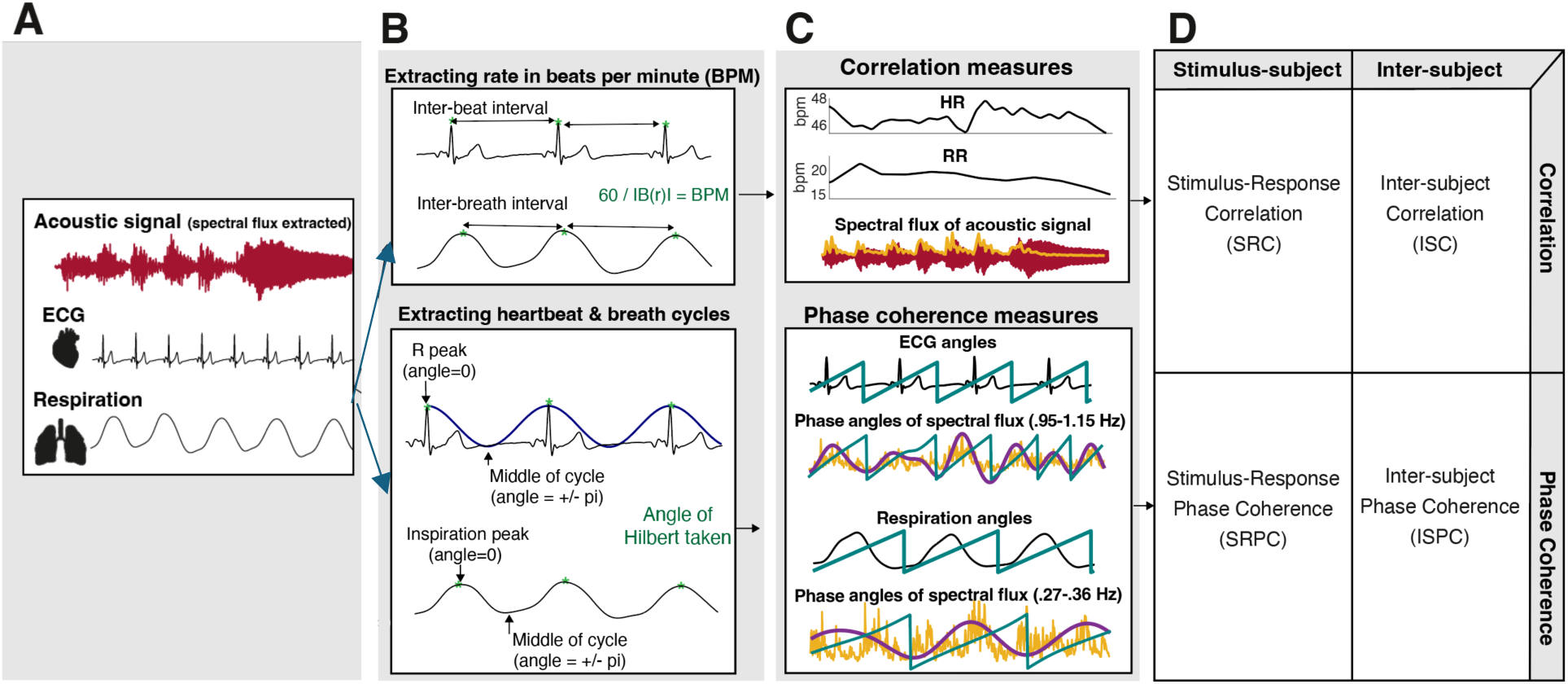
Outline of the experimental design and analysis pipelines. **Panel A** shows the study design: audiences were presented with music pieces in audio-visual (AV, orange box) and audio-only (AO, blue box) conditions, while heart (ECG), respiration, and acoustic signal of the music (maroon) were continuously recorded. **Panel B** shows how beats per minute (BPM) were extracted from the ECG and respiration signals, which were then used for correlational measures (upper panel). **Panel B** also shows that the angles of heartbeat and respiration cycles were extracted for phase coherence measures, where start of cycle began at peaks of ECG and respiration signals (peaks marked as green dots) (lower panel). **Panel C** shows the correlational measures (upper panel) and the phase-coherence measures (lower panel) we extracted from the continuous measures. For the correlational measures, we extracted heart and respiration rate (HR, RR, from cardiorespiratory signals) and spectral flux (from acoustic signal). For the phase coherence measures, we extracted the phase angle of ECG and respiration cycles (from cardiorespiratory signals), and angles of spectral flux band-pass filtered at the audience’s heart frequencies (0.951 - 1.159 Hz) and respiration frequencies (0.268-0.358 Hz). **Panel D** shows the extracted synchrony measures: stimulus-subject (vertical, left) and inter-subject (vertical, right) for correlation (horizontal, top) and phase coherence (horizontal, bottom) measures.

### Procedure

Participants were invited to attend one of two concerts. Electrocardiography (ECG) and respiration were collected using gelled self-adhesive Ag/AgCl electrodes and a respiration belt, respectively, and continuously measured during the concert at a 1000 Hz sampling rate. During the concert, audiences heard the three pieces both in AV and AO modalities. The order of modality was counterbalanced across the two concerts. After each piece, participants rated items such as engagement. We used a German word to reflect engagement as an absorbed state (see Introduction). Although it was translated as ‘absorption’ in a previous study^7^, it can also be used synonymously with engagement (see also^64^).

## Data Analysis

### Musical analysis: spectral flux and section boundaries

To model how humans respond to music, previous studies have extracted the envelope from the acoustic signal^4^. More recently, spectral flux has shown to be a better predictor of participants’ responses to an acoustic signal as it shares information of the acoustic envelope as well as note/speech onset information^23,24,65^. Therefore, the continuous spectral flux signal was obtained using the MIRToolbox^26^ in Matlab (2019b), with a frame size of 25ms (as is appropriate for short-term acoustic features^66^), with a 50% overlap (Figure 1b). To allow for comparison between conditions, we confirmed that spectral flux was similar across AV and AO performances, with Pearson correlations at *r* > 0.92 (see Supplementary Figure 1). To investigate the phase relationship between spectral flux against the cardiac and respiration measures fairly, we assessed spectral flux within the audience members’ average heart (1.01 Hz) and respiration (0.30 Hz) frequencies. This approach followed previous research that also selected spectral flux within specific frequencies (sub-bands) to better ascertain rhythm hierarchy levels^25,27^. Although it is possible in the MIRToolbox to specify these sub-bands, the default sub-bands were considered too wide for the current hypotheses of heart/respiration frequencies. Therefore, to obtain rhythmic levels that coincide with heartbeat and respiration rhythms, spectral flux was bandpass filtered at representative ranges of the interquartile range of all heart (0.951 - 1.159 Hz) and respiration (0.268-0.358 Hz) frequencies (see Supplementary Figure 2). The phase angle of filtered spectral flux was calculated from the real part of the Hilbert envelope using MATLAB’s *angle* function (Figure 1b, lower panel). The section boundaries in the musical pieces were identified either by a double bar line or end repeat bar line in the score, or by a change/repeat of thematic material (identified in the score by AMC, then confirmed by a music theorist, see Supplementary Table 1).

### Pre-processing of physiological data

Raw heart and respiration signals were pre-processed in MatLab 2019b. Missing data from the raw signals were first interpolated at the original sampling rate (all gaps in data were less than 60 ms). Data were cut per piece and further pre-processed using Fieldtrip^67^ and the *biosig* toolbox (http://biosig.sourceforge.net). Respiration data were low pass filtered at 2 Hz, ECG data were band-pass filtered between 0.6 and 20 Hz (Butterworth, 4^th^ order), and both demeaned. Peaks in ECG and respiration signals were extracted using, respectively, *nqrsdetect* function from biosignal and a custom-made script that located when the respiration signal exceeded a peak threshold. Computationally identified peaks were manually screened to ensure correct identification of R wave (part of the Q-, R- and S-wave [QRS] complex of ECG) and respiration peak locations. Any peaks that were not correctly identified were manually added, while falsely identified peaks were removed, for example, if a T wave in the ECG was accidentally identified as a R wave. In rare cases where participant data within a piece were too noisy to identify clear QRS and inspiration peaks, participant data for this piece were rejected from further analysis (ECG = 14%, respiration = 7%). The pre-processed heart (ECG) and respiration signals were then further analysed in two different ways to calculate synchrony: heart *rate* and respiration *rate* (BPM) were calculated for correlational synchrony, while *phase angle* of ECG (phase of heartbeat cycle) and respiration (phase of inspiration/exhalation cycle) were calculated for phase synchrony (more details outlined below, see Figure 1A-B).

### Synchrony analyses

Four continuous synchrony measures were extracted: stimulus-response correlation (SRC), inter-subject correlation (ISC), stimulus-responses phase coherence (SRPC), and inter-subject phase coherence (ISPC) for each ECG and respiration (see Fig. 1D).

### Correlational synchrony: ISC and SRC

Correlational synchrony measures (SRC, ISC) were calculated using heart rate (HR) and respiration rate (RR). To do so, the heart (ECG) and respiration signals converted into rate (beats per minute, BPM) by first obtaining the differential timing between peaks, i.e., inter-beat intervals (IBI) for ECG, and inter-breath intervals-(IBrI) for respiration (see Figure 1B). IB(r)Is were then converted to BPM and interpolated at the original sampling rate to obtain instantaneous HR and RR, which were downsampled to 20 Hz^68^. SRC and ISC were then calculated across sliding windows. Although ISC and SRC measures have previously been calculated within 5 second sliding-windows for EEG^4,12^, peripheral responses of heart and respiration are slower, where a typical evoked response can be up to 10 seconds^14,16,69^. Therefore, we chose a 10 second sliding-time window with a one second overlap. For SRC, each participants’ HR and RR signal was correlated with the extracted spectral flux. For both HR and RR signals, *p* × *t* matrices were created, where *p* was the HR or RR signal for each participant over *t* timepoints. Signals were correlated in a pairwise fashion, i.e., between all possible participant pairs within one concert. Values underwent Fisher-Z transformation, were averaged, and then transformed back to r values (inverse Fisher-Z).

### Phase coherence synchrony: ISPC and SRPC

Phase coherence measures were used to assess whether heartbeats and breaths aligned with the spectral flux rhythms in music. For this, heart and respiration signals were transformed into the phase domain. Here, we assessed changes of actual heartbeat and breath cycles that vary over time. To account for these variations, phase was calculated from cycle peaks, see Figure 1B, green dots marking the R peaks in the heart signals and inspiration peaks in the respiration signals. For the phase angle of the heartbeat, a value of 0 denoted the R wave (peak of heartbeat cycle) and ±***π*** the midpoint of the cycle (see Figure 1B). For the phase angle of the respiration cycle, 0 denoted the inspiration peak, while ±***π*** represents the midpoint of the cycle or maximum exhalation (see Figure 1B). To obtain such values, a continuous sinusoidal wave was fitted to each IB(r)I of detrended and normalised ECG and respiration signals based on^70^, using the following equation:

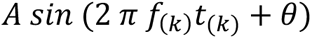

where *A* is the mean peak amplitude (i.e., average amplitude of signal at time points of QRS/respiration peak), *f*_(*k*)_ is the frequency calculated from IBI (i.e., peak_k+1_-peak_k_), converted to Hz 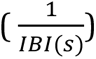. Phase (*θ*) was optimised so the peak of the sine wave corresponded to the ECG/respiration peak. Next, the phase angle was calculated from the real part of the Hilbert using MATLAB’s *angle* function.

SRPC of heartbeats was calculated by assessing coherence between the phase angles of heartbeat cycles with the spectral flux phase angles corresponding to heart (bandpass filtered at 0.951 - 1.159 Hz), while SRPC of respiration was calculated by assessing coherence between phase angles of respiration cycles with spectral flux phase angles corresponding to respiration (bandpass filtered 0.268-0.358 Hz) frequencies (see Figure 1C). Coherence was calculated based on the following formula based on^71^:

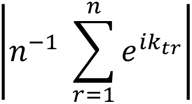

where *n* is the number of phase signals, *e^ik^* is Eulers formula (i.e., the complex polar representation of phase angle *k* for signal *r* at time point *t*). ISPC was assessed in a pairwise fashion between heartbeat cycle and respiration cycle phase angles of all possible participant pairs within one concert using the same coherence formula as above.

### Adjusting for lags

To account for lags in responses to the stimuli for SRC and SRPC, we adjusted data with a constant lag to optimally align the stimulus and the corresponding responses. This lag was informed by previous literature that added lags to account for slight delays in stimulus responses^4,72–74^. To do this, we first assessed lags on an individual basis. Lags between a stimulus and each participant’s response were calculated within the first 10 seconds after stimulus onset as this initial onset likely evokes the most reliable responses. We calculated SRC and SRPC at lags up to each individual’s full cardiac and respiration cycle (based on^72^); that is, their mean inter-beat-interval for heart measures (on average: 990 ms) or inter-breath interval for respiration measures (on average: 3400 ms), as it might take up to one heartbeat/respiration cycle to begin responding to a stimulus^14^. The positive correlations and phase coherence values were obtained at the optimal lag^73^. We then averaged the individual optimum lags^74^. This average lag was applied for every participant (i.e., each participant had the same lag for each stimulus-response synchrony measure). For correlation values, the average optimal lags for HR and RR were 579 ms and 1573 ms, respectively. For phase coherence values, the average optimal lag for heart and respiration phase cycles were 500 ms and 1820 ms, respectively. These lags were applied to all corresponding heart and respiration stimulus-response pairings for the correlation and phase coherence. Therefore, for each stimulus-response synchrony measure, all participants had the same lag.

### Synchrony across (time-averaged) and within (time-resolved) pieces

Continuous synchrony measures for correlational measures (SRC-HR, ISC-HR, SRC-RR, ISC-RR)) and phase measures (SRPC and ISPC of heartbeats, and SRPC and ISPC for respiration phase) were calculated in a time-averaged or time-resolved fashion to test the different hypotheses. For the first two hypotheses, we calculated time-averaged synchrony values. Rather than averaging synchrony across the three music pieces - as three musical pieces were thought to yield too few observations - we obtained time-averaged synchrony per piece section. The Bach piece was split into seven sections, the Beethoven piece was split into nine sections, and the Messiaen piece was split into nine sections (for details see Supplementary Table 1). Synchrony was then averaged per piece section, yielding 25 observations per participant, per condition, and per synchrony measure. Averaging across musical sections was to account for naturalistic grouping as defined by the musical composition. As an additional check, we also calculated average synchrony within 30 second bins, thereby controlling for length and section changes (yielding 53 observations per participant, per condition, per synchrony measure, see Supplementary Materials), and ran identical statistics. To address the third hypothesis, we calculated time-resolved synchrony values by cutting epochs ±15 seconds relative to section boundary onsets, to capture event-related respiration and heart responses^14,69^, as well as any anticipatory effects at musical events^75^.

### Significance-testing

Regarding the first hypothesis, we tested the assumption that responses time-locked to a stimulus should evoke similar responses across participants^28^. Therefore, we created time-‘unlocked’ data by circular shifting ECG and respiration signals. This was done 1000 times and time-averaged synchrony measures were calculated as described above. For each of the permutated data sets, we calculated a *t*-test statistic, creating a null distribution of permutation values. By assessing how many of these permutation values were higher than or equal to the statistic of the true (time-locked) data, we determined the chance of true synchrony (see Supplementary Figure 3). Values were corrected for multiple comparisons using false discovery rate (FDR^76^).

### AV versus AO: time-averaged synchrony

For the second hypothesis, time-averaged synchrony measures were compared between AV and AO. Self-reported engagement was also compared between these conditions. Linear mixed models (LMMs) with a fixed effect of modality were constructed for each of the synchrony measures as the outcome variables. As the experimental design meant that data were clustered, the following random effects were added to account for this non-independence: random intercepts for concert, piece, and participants, where participants were nested within concerts, while participant and piece were considered crossed effects. A random slope for participants was also included. This random-effect structure represents maximal models^77,78^. If these maximal models generated convergence and/or singularity fit errors, models were simplified following recommendations^77,79^. As output from models generating errors should not to be trusted^77^, we report simplified, error-free models in the results. Maximal model outputs can be nonetheless found in Supplementary Materials, while all LMMS can be found in accompanying code on OSF (see link/Footnote (1) above). LMMs were run using *lmer* from the *lme4* packages^80,81^. Significance values and effect sizes were obtained from the *tab_model* function from *sjPlot* package^82^. As a recommended sanity check, we also ran conventional *t*-tests alongside LMMs^83^. This was done by a one sample *t*-test (or Wilcoxon if data were not normally distributed), run on the difference between AO and AV values.

### AV versus AO: epoched

To test the third hypothesis, synchrony at section boundaries was assessed between modalities across time windows. Synchrony values in the 30-second epochs centred around section boundaries were split into smaller time windows^69,84^, yielding five 6-second time windows. For each synchrony measure, linear mixed models (LMMs) with fixed effects of modality and time window were constructed. We added random intercepts for concert, piece, and participants (participants nested within concerts; participant and piece as crossed effects) and a random slope for participants. As above, models generating errors were simplified; error-free models are reported (full models in Supplementary Materials). Estimated marginal means (using *emmeans* package^85^) were used to check pairwise comparisons with Bonferroni adjustments. As we predicted that synchrony would occur at section boundaries, we expected the time window centred at the section boundary to have significantly higher synchrony values than time windows prior to and after a section boundary. In case synchrony was high, we investigated what responses were becoming synchronized. Thus, HR and RR data at section boundaries were taken (i.e., HR and RR as in Figure 1, Panel B). Two LMMs were constructed, each for aggregated HR and RR values, with fixed effects of time window and random effects of participant id, piece, and concert, as above.

## 3. Results

### Significance of synchrony measures

In assessing the significance of synchrony measures using a permutation approach and FDR correction, we found that SRC-HR (*p* < .001), ISC-HR (*p* = .048), and ISC-RR (*p* = .048) showed significant, above-chance synchrony (see Supplementary Figure 3). SRC-RR (*p* = .408), SRPC for heartbeats (*p* = .412) and respiration (*p* = .594), and ISPC for heartbeats (*p* = .594) and respiration (*p* = .594) were not above chance level. When synchrony values were averaged in 30 second bins, significant results were replicated for SRC-HR (*p* < .001), but not for ISC-RR (*p* = .051) and ISC-HR (*p* = .065). We therefore carried through SRC-HR, and (tentatively) ISC-RR and ISC-HR measures for further modality comparisons.

### Time-averaged synchrony between modalities

Figure 2 shows that compared to the AO condition, the AV condition evoked generally higher SRC-HR, ISC-HR, and ISC-RR, but only for ISC-HR was this modality difference statistically significant (see Table 1). This effect for ISC-HR occurred in both error-free and full models (see Supplementary Table 6). When checking conventional *t*-/Wilcoxon tests (depending on normality), results were similar to LMM results, though ISC-HR was no longer significant (SRC-HR, *t*(21) = 1.176, *p* = 0.253; ISC-HR, *t*(21) = 1.890, *p* = .073; ISC-RR, *t*(22) = .447, *p* = .659). Although self-reports of engagement were also higher in AV (see Figure 2), this difference was not significant in the LMM (ß = 0.33, 95% CI [-0.10-0.76], *p* = .129, R^2^_(fixed)_ = 0.007), nor in the traditional *t*-test (*t*(24) = 1.901, *p* = 0.069). In checking the behavioural relevance of synchrony as an engagement index, ISC-HR was not significantly correlated (rho = -0.093, *p* = .295) nor a significant predictor of self-reported engagement (ß = -0.001, 95% CI [-0.003-0.001], *p* = .464, R^2^_(fixed)_ = 0.005).

**Figure 2.**
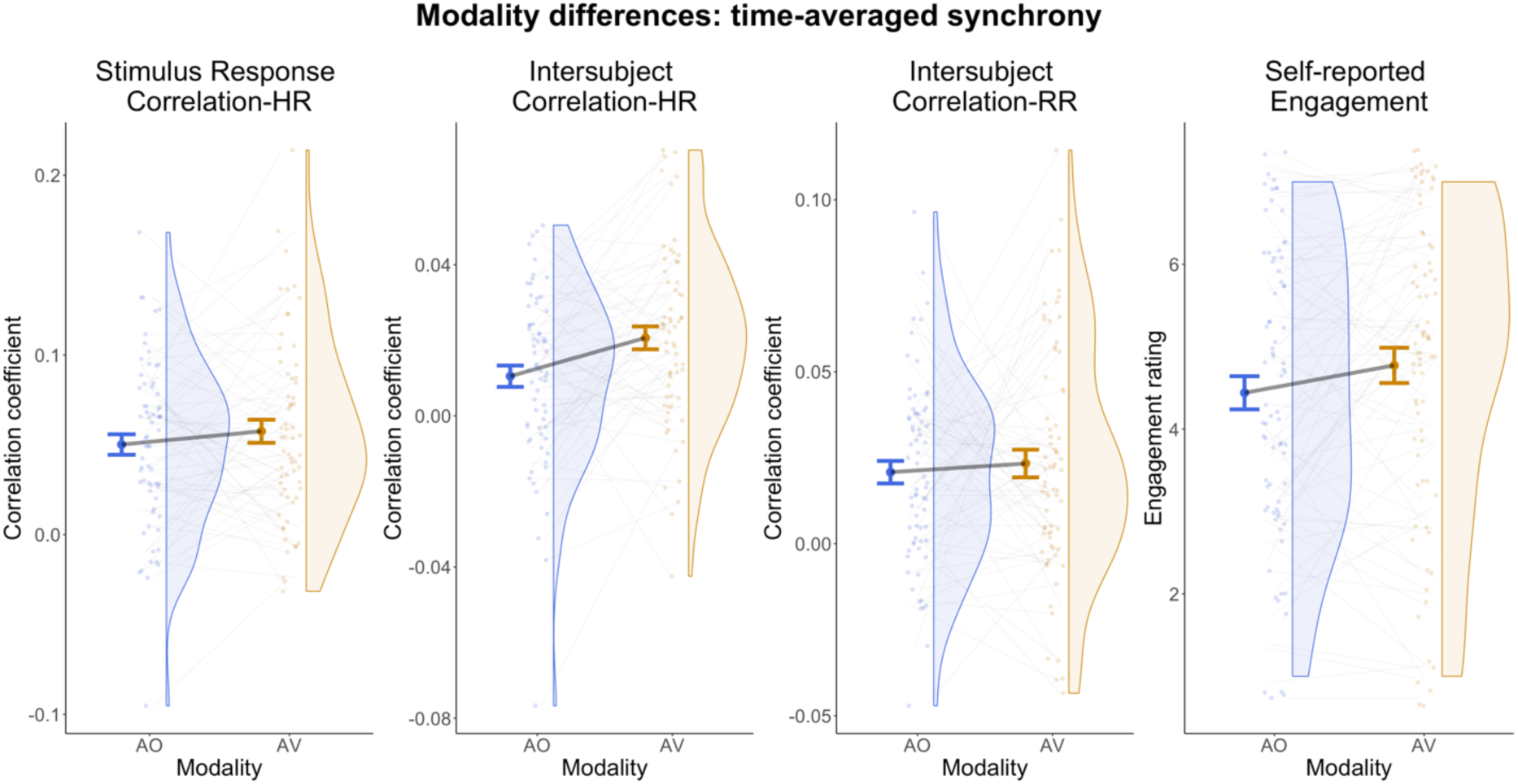
Raincloud plots showing modality differences of audio-only (blue) and audio-visual (orange) conditions for time-averaged synchrony measures of SRC-HR (far right), ISC-HR (right), and ISC-RR (left) as well as self-reported engagement (far left). Smaller points represent each data point, while the larger point represents the mean; error bars represent standard error of the mean. The shape of the distribution is shown by the half violin.

**Table 1.** Linear mixed models comparing synchrony between modalities.

| Predictors | SRC-HR |  |  | ISC-HR |  |  | ISC-RR |  |  |
| --- | --- | --- | --- | --- | --- | --- | --- | --- | --- |
|  | Estimates | CI | p | Estimates | CI | p | Estimates | CI | p |
| (Intercept) | 0.050 | 0.022 – 0.079 | **0.001** | 0.012 | 0.002 – 0.022 | **0.020** | 0.021 | 0.010 – 0.032 | **<0.001** |
| mod [AV] | 0.009 | -0.006 – 0.024 | 0.235 | 0.010 | 0.001 – 0.019 | **0.028** | 0.002 | -0.007 – 0.012 | 0.633 |
| **Random Effects** |  |  |  |  |  |  |  |  |  |
| $\sigma^2$ | 0.01 | | | 0.00 | | | 0.01 | | |
| $\tau_{00}$ | 0.00<sub>mac:conc</sub> | | | 0.00<sub>piece</sub> | | | 0.00<sub>mac:conc</sub> | | |
|  | 0.00<sub>piece</sub> |  |  |  |  |  | 0.00<sub>piece</sub> |  |  |
| $\tau_{11}$ | 0.00<sub>mac1.modAO</sub> | | | 0.00<sub>mac1.modAO</sub> | | | | | |
|  | 0.00<sub>mac2.modAV</sub> |  |  | 0.00<sub>mac2.modAV</sub> |  |  |  |  |  |
| $\varrho_{01}$ | | | | | | | | | |
| $\varrho_{01}$ | | | | | | | | | |
| ICC | 0.08 |  |  | 0.05 |  |  | 0.01 |  |  |
| N | 3<sub>piece</sub> |  |  | 3<sub>piece</sub> |  |  | 3<sub>piece</sub> |  |  |
|  | 16<sub>mac</sub> |  |  | 16<sub>mac</sub> |  |  | 16<sub>mac</sub> |  |  |
|  | 2<sub>conc</sub> |  |  |  |  |  | 2<sub>conc</sub> |  |  |
| Observations | 1075 |  |  | 1075 |  |  | 1152 |  |  |
| Marginal $R^2$ / Conditional $R^2$ | 0.002 / 0.079 | | | 0.007 / 0.056 | | | 0.000 / 0.011 | | |

### Synchrony increases at section boundaries, depending on modality

LMMs showed that in time-resolved epochs centred around section boundaries, SRC-HR and ISC-HR were significantly higher in the AV condition, while ISC-RR was higher in AO (see Figure 3 and Table 2, see Supplementary Table 7 and code for maximal models, which replicated these effects).

**Figure 3.**
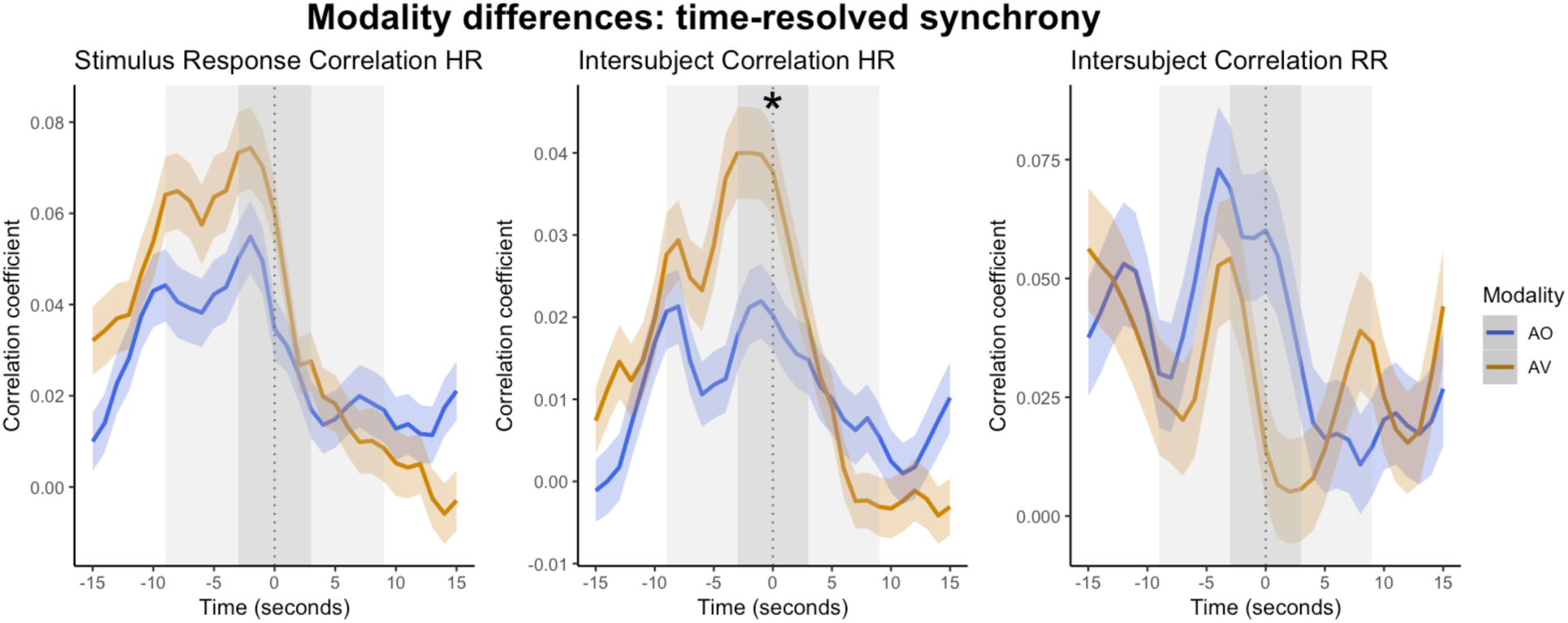
Synchrony values at 15 seconds prior to and after section boundaries for stimulus correlation of HR (left), inter-subject correlation of HR (middle) and inter-subject correlation of RR (right). The zero point represents the section boundary. Thick lines represent the average, and the ribbon represents the standard error of the mean for audio-only (AO, blue) and audio-visual (AV, orange) conditions. * represents a significant modality effect within that time window.

**Table 2.**
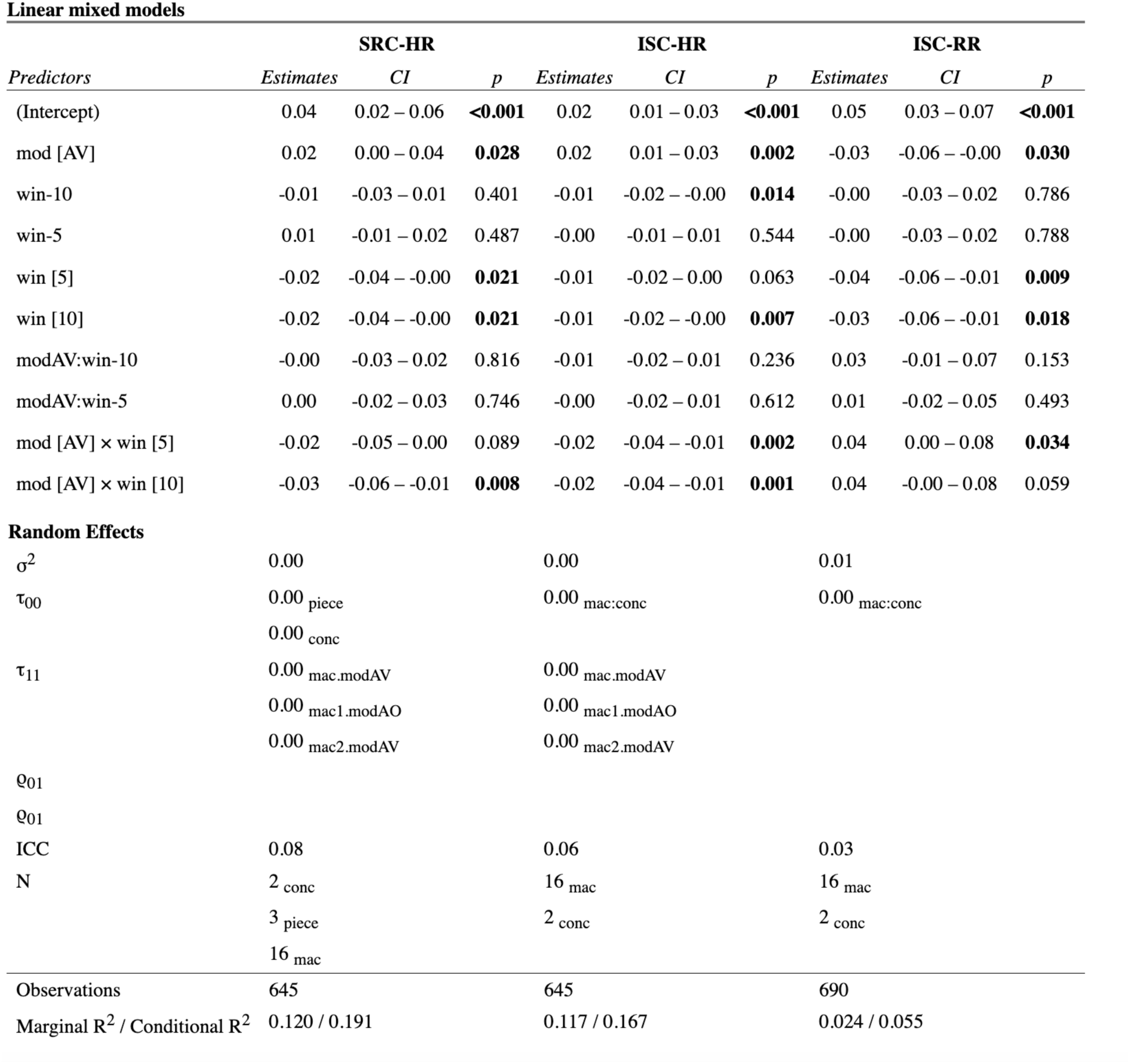
Linear mixed-effects models for synchrony with predictors of time window and modality.

LMMs additionally showed that time windows predicted synchrony measures at section boundaries. Synchrony measures were significantly higher in the time window centred at the section boundary compared to time windows before (10 seconds before: ISC-HR), and after (5 seconds after: SRC-HR, ISC-RR; 10 seconds after SRC-HR, ISC-HR, and ISC-RR) (see Figure 3 and Table 2).

In assessing the interaction between time window and modality, Figure 3 shows that synchrony was higher before and at section boundaries (window 0) compared to 5 and 10 seconds after section boundary in the AV condition for SRC-HR and ISC-HR. This latter effect was further confirmed by LMMs and by significant pairwise comparisons (Bonferroni corrected) for ISC-HR (Table 3). ISC-RR was higher in AV just after (5 second) section boundary, but this effect was not significant in Bonferroni corrected pairwise comparisons (Table 3).

**Table 3.**
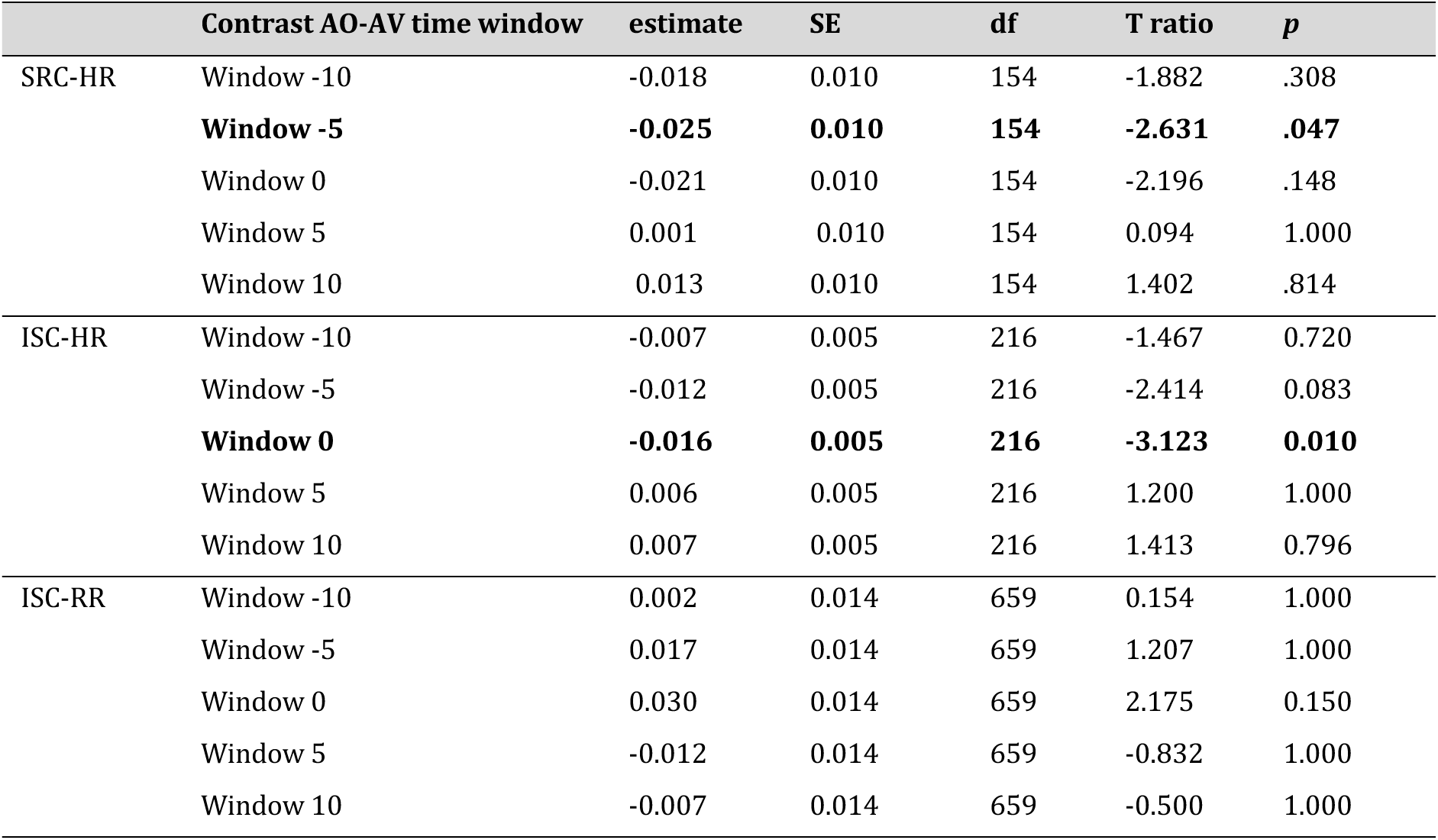
Pairwise comparison (Bonferroni corrected) between modality (AO-AV) in each time window.

Upon checking actual HR and RR responses at these time points^16^, this synchrony corresponds to typical orienting responses (see Figure 4). HR follows a deceleration-acceleration pattern, where the estimate in HR in window-10 and window 10 is significantly higher than window 0 (intercept, see Table 4 and Supplementary Table 8). Although RR decelerated and accelerated similarly to HR (see Figure 4), these were not significant across time windows.

**Figure 4.**
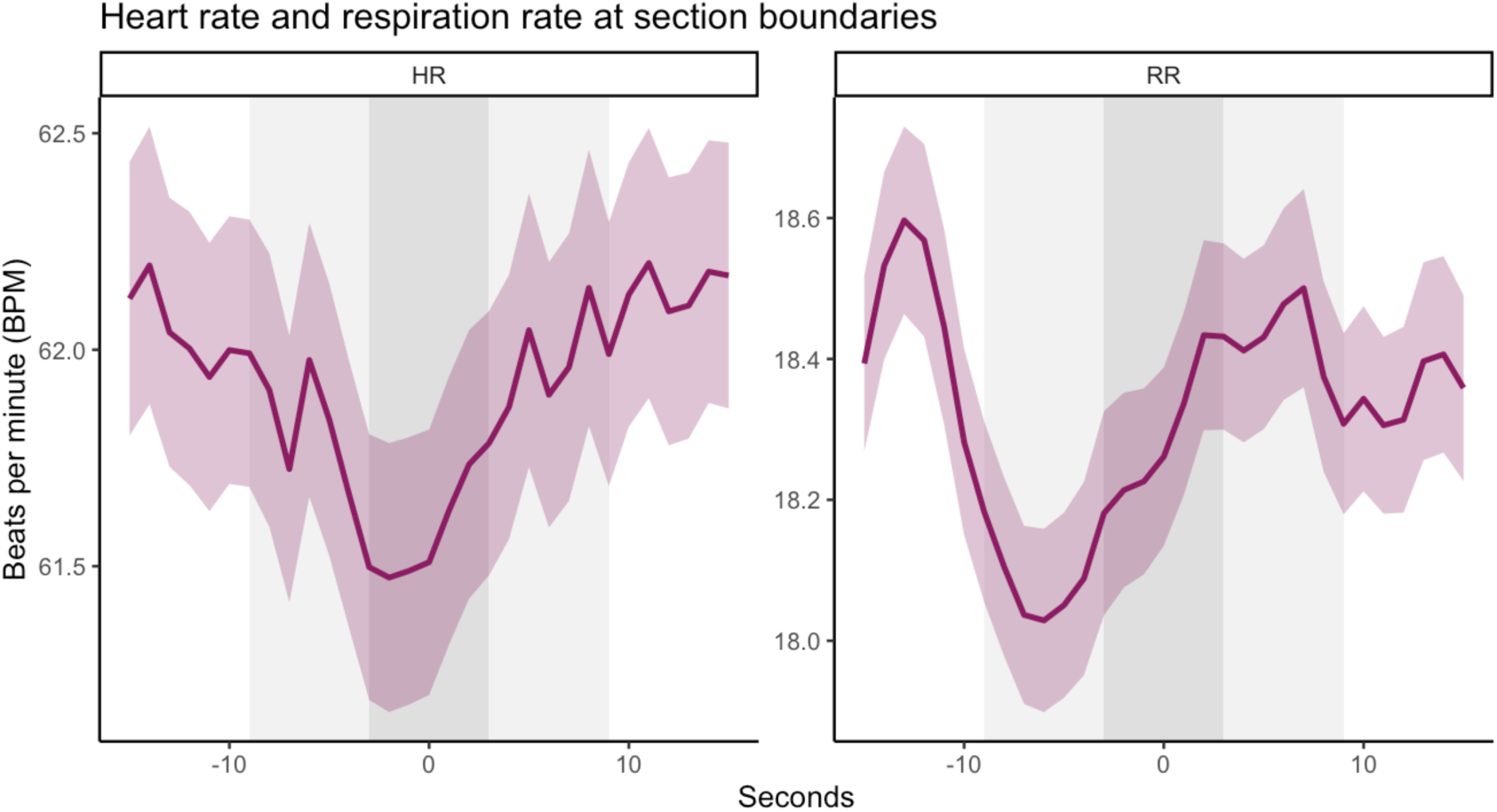
Heart rate (HR, left) and respiration rate (RR, right) BPM values at 15 seconds prior to and after section boundaries. The zero point represents the section boundary. Thick lines denote the average BPM, while ribbons around the average line represent the standard error.

**Table 4.**
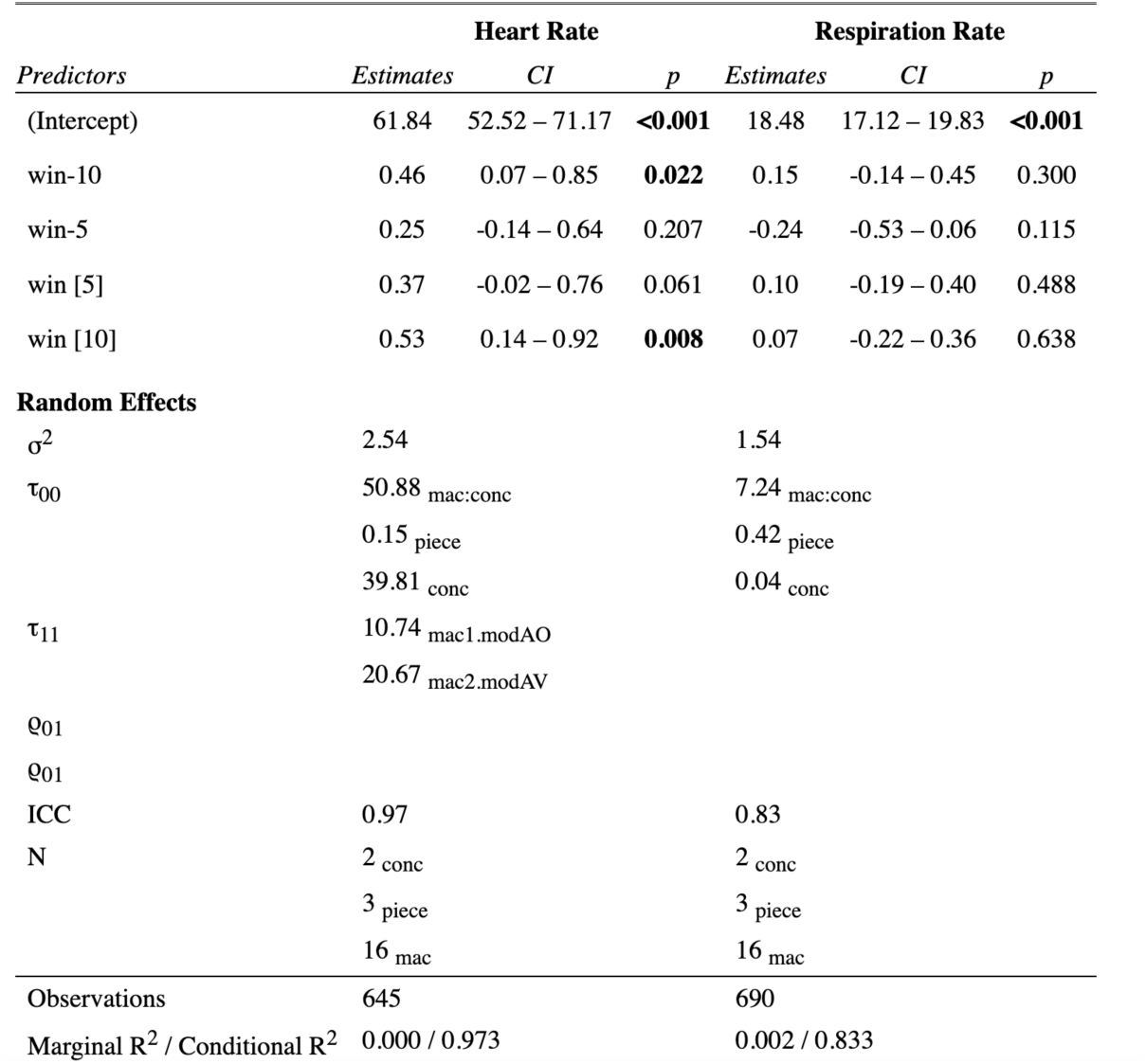
Linear mixed models for raw heart and respiration rate at epochs centred around section boundaries.

## 4. Discussion

An important aspect of (live) musical experiences is the listeners’ engagement. Despite the methodological challenges in assessing such musical experiences, dynamic engagement is increasingly assessed with neurophysiological synchrony measures. By extending these techniques to a naturalistic setting in a concert hall, the current study pushed the boundaries in how to measure and analyse musical experiences. Importantly, we also compared approaches to gauge the robustness of these cardiorespiratory synchrony analyses. As engagement might be enhanced by seeing musicians perform^8,10^, participants were presented with AO and AV performances of Western classical music in a concert setting while cardiorespiratory measures were continuously measured. We show that heart rate (i.e., *speed* of heartbeats in BPM) correlated with spectral flux of music above chance. Additionally, respiration rate and (tentatively) heart rate correlated across audience members above chance. Musical rhythms did not align with heartbeat rhythms nor breathing rhythms above chance. We also find that compared to the AO condition, AV music performances evoked significantly higher ISC-HR; this effect was more robust on a time-resolved, compared to time-averaged, level.

### Audience heart rate and respiration rate correlation

HR correlated with spectral flux (SRC-HR) above chance level, indicating that the acceleration/deceleration of audiences’ HR correlated with increases/decreases of spectral flux in the music. ISC-HR and ISC-RR measures were also above chance level, indicating that acceleration/deceleration of HR and RR correlated across audience members experiencing music simultaneously, supporting previous research showing that ISC of HR^18,46^ and EEG^4^ is evoked when participants simultaneously experience the same auditory stimuli. We note, however, that although ISC-HR was significant in the main analysis of time-averaged synchrony within musical sections (see Supplementary Table 1), when ISC-HR was time-averaged across 30-second bins this measure did not meet the significance threshold after correcting for multiple comparisons. Therefore, it might be important to assess synchrony related to musical structure and on a time-resolved level; see below discussion on ‘Time-resolved synchrony’. That ISC-RR was above chance level somewhat contrasts findings of Madsen and Parra^46^, who found ISC-RR not to be significant. One explanation could be due to stimuli differences. Here, we used music stimuli, whereas Madsen and Parra used instructional video (speech). More importantly, the setting of a concert hall might have increased the engagement of the listeners compared to the laboratory setting. Our findings suggest that HR and RR (i.e., speed of heartbeats/breaths) may become synchronized across groups of people experiencing auditory stimuli (like music) simultaneously, but may be context dependent.

### Heartbeat and breathing cycles do not rhythmically align to music

Both musical rhythm–heartbeat phase synchrony and musical rhythm–breath phase synchrony (SRPC measures) were not significantly above chance level. In other words, musical rhythms aligned with neither audiences’ breathing cycles nor their heartbeat cycles. Although this lack of phase synchrony was expected for heart rhythms^51–53^, this contrasts previous findings by Etzel et al^49^ and Haas et al^50^ that breathing aligns with musical beats. There are a few potential reasons for this discrepancy between results. First, Etzel et al. and Haas et al. compared respiratory signals to the beat of music, while the current study assesses spectral flux filtered at respiration frequencies. Although our method is therefore not comparable with this previous research, we nonetheless suggest that our method is more generalisable for future naturalistic research. It seemed like the previous research chose music to roughly correspond to healthy breathing frequencies (see Haas et al, Table 1, Figure 1 and 2; Etzel et al. Table 1). However, much music might not correspond to such specific frequencies. Indeed, the musical choice for the current study was motivated by the need to present a typical musical program, where the beat did not fall into the range of healthy heartbeats/breathing (see Supplementary Table 1). It therefore seemed problematic to assess synchrony of the actual beat with heartbeats/breaths. Assessing musical features that strongly correlate with beat onsets (spectral flux^24^) nonetheless yields a somewhat comparable approach. Additionally, extracting the energy at the natural frequency of heartbeats/breathing allows assessing music that has beats falling outside of the natural breathing/heartbeat range, providing a useful measure for future research. A second explanation for not replicating musical beat-respiration synchrony, is that previous studies used music with relatively regular rhythmic and metric structures^49,50^, while the stimuli in the current study included one contemporary piece with ambiguous rhythms. However, there were no respiration synchrony differences between pieces with regular (Bach, Beethoven) and irregular (Messiaen) metric structures (see Supplementary Table 3). A third explanation could be that respiration is more likely to align with beats in laboratory experiments as the attention (per instruction) is mainly on music. Beyond complexities in musical structure, a real-world setting presents several additional factors, such as the visual and social aspects^10,86^, where attention may fluctuate a lot more between the music and other features. Thus, although neural^72,74^ and respiratory^49,50^ rhythms may align to auditory beats in a laboratory setting, the current results suggest that this may not occur to music with broader tempi in a music concert setting, at least for this choice of Western Classical music.

Heartbeats and breaths likewise did not significantly align in phase between participants. One explanation could be the lack of social interaction, which is a crucial aspect in higher-order phase coupling^40^. Indeed, phase synchrony between participants has previously been found in dancing^87,88^ and musical ensembles ^38,39,89,90^ where there is a clear shared goal of a coordinated performance (see^40,41^). Typical Western classical concert etiquette calls for enhanced focus on the performance, thus reducing body movements and interaction between audience members^10^. Indeed, ratings for ‘urge to move to the music’ (see^7^) were heavily skewed to low ratings; even testing significance of phase coherence in audience members who only had high ‘urge to move’ ratings did not yield significant results (see Supplementary Table 5). Additionally, videos showed that the audience members respected this etiquette and generally did not move too much^7^. Therefore, we suggest that phase synchrony – typically seen in interactive settings^40^ – may not occur in non-interactive settings.

### Time-averaged synchrony may not relate to naturalistic engagement

Next, synchrony measures were compared between AV and AO conditions. ISC-HR was significantly greater in AV performances. However, this result was not shown to be robust with a sanity check using conventional *t*-test. Although our data are more appropriate for LMMs due to naturalistic paradigm and physiological data^78^, the fact that the significance is not replicated in a simpler *t*-test thus allows only cautious interpretation^83^. Another reason to be cautious is that – contrary to what was expected – ISC-HR was not significantly related to engagement that was self-reported after each piece. The current findings contrast findings from Ki et al.^31^ who found that neural ISC was stronger in attend than non-attend conditions, especially in audiovisual stimuli. The current finding also contrasts studies showing that HR^18^ and neural^19^ synchrony can reflect engagement. One explanation for the discrepancy in results is that these previous studies had explicit attend/non-attend conditions, where participants were instructed to count backwards in the non-attend conditions. Here, participants experienced concert performances naturally with no formal instruction. This suggests that while ISC (of HR) could be a marker in clearly defined engagement conditions, it might not necessarily be generalisable to less controlled settings. Additionally, inter- and intra-individual differences may influence synchrony and engagement^55,91,92^, and could be a potential venture for future studies to explore individual versus group synchrony. Overall, we show time-averaged synchrony does not reliably reflect overall self-reported engagement measurements. With this in mind, we anticipated results for time-resolved synchrony might yield clearer results.

### Time-resolved synchrony reveals synchronized heart rate orienting response

Engagement with music typically fluctuates across time^4,12,54^. Thus, we assessed synchrony changes between modalities on a time-resolved scale. We ‘zoomed in’ on synchronized responses at salient moments in the music, here defined as section boundaries in music (see Supplementary Table 1), which are important structural locations in music. The current results show that both ISC-HR and SRC-HR increased at such section boundaries, supporting and extending previous work on ISC^12,16^ to SRC. Although we did not have time-resolved self-reports to assess the relation of synchrony to engagement, we assessed the continuous HR and RR. We show that synchronized responses reflected significant deceleration-acceleration patterns in HR and RR. Such patterns have been long associated with a typical orienting response, which marks an increase in attentiveness^93^ to anticipate and perceive events^14,94–96^. Thus, such increases of cardiorespiratory synchrony at section boundaries suggest increased engagement with music at these time points, related to auditory cues indicating a section is ending, such as a cadential ending (i.e., harmonic closure in Bach and Beethoven) and a slight slowing down or change in tempo and/or material (see Supplementary Table 1). This physiological synchrony at salient music events suggests a bottom-up capturing of engagement during a shared musical experience^40^. We acknowledge that the definition of salient moments to explore engagement are limited in the current study to section boundaries. However, humans also respond to segmentation not only at section boundaries^4,12,61,97^, but also at smaller phrasal^98^ and larger, movement^68^ levels. Thus, the current work provides a starting point for exploring engagement to salient moments; future work could compare ‘non’ salient moments, transitions, and segments at lower and higher hierarchical musical levels.

Importantly, Figure 3 shows that ISC-HR significantly increased at section boundaries more so in the AV condition. This increase suggests that visual information further increased engagement at section boundaries. Indeed, previous performance studies have found that certain gestures cue and anticipate structurally important locations, such as leaning forward towards a cadence(harmonic closure)^99^, increased movement amplitude^100^, or finishing ‘flourish’ gestures^101^ as well as deep breaths and sweeping motions prior to onset of new phrases^102^. Indeed, videos show that the pianist tended to lean forward at section boundaries (though see limitations regarding analysis of visual features). Therefore, a combination of audio and visual aspects likely increases engagement with music at a local level, as shown most robustly with ISC-HR.

## 5. Limitations

Although we show greater synchrony in the AV (compared to AO) condition, it is unclear whether this synchrony increase might simply be related to increased information processing of both visual and audio information, compared to simply audio information. For example, RR of audiences did not synchronise to the spectral flux of the music (insignificant SRC-RR result), but RR did significantly synchronise across audience members (ISC-RR). This suggests that the audiences’ RR synchrony was rather driven by something else than the acoustic signal, perhaps the musician movements (see^7^). Indeed, we saw that SRC-RR was more significant in the AV than the AO condition (see Supplementary Table 4). As it was not possible to extract reliable visual features in the current study due to the camera location in the concert hall (see angle in Figure 1A), future work would therefore need to assess responses not only to auditory features, but also visual features, such as facial expressions^103^, quantity of motion, and speed of movements^104^ to shed more light on how much variance in the audience experience can be explained by auditory and by visual features. Additionally, we did not test a visual only condition which would allow us to disentangle the difference between added information processing and actual engagement. Finally, we acknowledge that aspects of ‘liveness’ may have likewise contributed to the current results. For example, previous evidence suggested that musicians have different perceived expressiveness when performing with no audience (rehearsal condition) compared to performing in front of an audience (recital condition^105^).

However, as the literature is mixed as to whether liveness influences emotional audience experience^106^ (for further discussion see^7,107^), but the literature on audio-visual influences on music appreciation is relatively robust^9^, it seems more plausible that the current results are explained by the modality difference rather than liveness.

## 6. Conclusion

The current study pushed the boundaries of how to measure and analyse music listening experiences. One crucial aspect of such a musical experience is the listener’s engagement, which can be assessed by measuring neurophysiological synchrony. As engagement could be affected by modality, we explored engagement differences between audio-visual and audio-only music performances in a Western classical concert setting, using cardiorespiratory synchrony as a measure of engagement. We extended studies investigating neural synchrony^4,17^, by comparing the robustness of different analyses to calculate peripheral physiological synchrony^18,46,54^ in music concert settings^2,15,16^. We showed that seeing musicians perform can heighten an audience’s engagement, though this might be context-dependent, and is best assessed by time-resolved (rather than time-averaged) inter-subject correlation of HR (i.e., synchrony of heartbeat speed). Rhythmic alignment (phase synchrony) of heartbeats/respiration with other audience members/music) were below chance level. However, questions arose whether phase coherence might become more relevant in contexts where listeners can move and interact, for example, at pop and rock concerts. More studies, involving a wider range of genres and contexts are needed to further understand synchrony dynamics in real-world music contexts. Importantly, the current study highlights the advantages of timeseries analysis in quantifying dynamic engagement, which could be applied not only to data collected during concert music listening, but in different naturalistic settings (e.g., open air concerts).

## Acknowledgements

The authors would like to thank Lea Fink, our music theorist who advised on the musical sections. Thanks also go to Klaus Frieler and Örjan De Manzano for discussions about statistics. Many thanks also to the ArtLab team, who assisted in concert preparations.

## Footnotes

1 Please note this repository is currently private and anonymous while this manuscript is under review; it will be made public upon acceptance of this manuscript.

