## Supplementary Material for "Audio-visual concert performances synchronize an audience’s heart rates"

### Supplementary Materials

**Supplementary Table 1:** Outline of the sections. For Bach's Prelude and Fugue, the sections were clear with double bar lines, repeat bar lines, or a change in thematic material, tempo and/or harmony. For Beethoven's Sonata, section changes occurred mainly with changing material and/or double bar line. For Messiaen, section changes occurred with changes of tempo (indicated by the composer) and/or double bar lines. Supplementary Table 1 presents a brief description of each section.

| Piece | Section | Bar | Metre | Tempo | | Description |
|---|---|---|---|---|---|---|
| | | | | M | SD | |
| Bach | 1 | 1 | 12/8 | 88 | 5 | First section of prelude, main theme of ascending semiquaver and triplet pattern followed by descending quavers. |
| | 2 | 17 | 12/8 | 86 | 4 | Repeat of first section of prelude - exact same music as in previous section. |
| | 3 | 33 | 12/8 | 84 | 4 | Second section: development of semiquaver-triplet-quaver theme. |
| | 4 | 57 | 12/8 | 86 | 3 | Repetition of introductory material (i.e., ascending and descending patterns as in bars 1-17) in the second section. |
| | 5 | 73 | 12/8 | 84 | 4 | Repeat the second section. |
| | 6 | 97 | 12/8 | 82 | 11 | Repeat of introductory material in the second section. |
| | 7 | 113 | 2/2 | 88 | 9 | Fugue in four voices. Musical material has a natural development of subject in a clear 2/2 pulse. |
| Beethoven | 1 | 1 | 6/8 | 48 | 3 | First section. Quiet, relatively clear quaver pulse. |
| | 2 | 9 | 6/8 | 54 | 5 | Second section. Bass has semi quaver Alberti bass. |
| | 3 | 17 | 6/8 | 50 | 7 | Dynamics changing loud, forte, reinforced chords. |
| | 4 | 26 | 6/8 | 51 | 5 | Quaver pulse in left hand, nice quiet melody in right hand, which gets louder over time. |

| | | | | | | |
|---|---|---|---|---|---|---|
| | 5 | 30 | 6/8 | 58 | 9 | Triplet quaver pulse in left hand, with low pit low pitch right hand, followed by minor, followed by high-pitched demisemiquavers in right hand. |
| | 6 | 44 | 6/8 | 58 | 5 | Repetition of the first section material. |
| | 7 | 56 | 6/8 | 53 | 6 | Repetition of section 3 material. |
| | 8 | 65 | 6/8 | 50 | 8 | Quaver pulse LH, RH semidemi triplets, which speed up to hemidemisemiquavers, start quieter and lower in register, increase in pitch and dynamic. |
| | 9 | 76 | 6/8 | 58 | 10 | Closing section. Quaver pulse mostly with semitone intervals |
| Messiaen | 1 | 1 | | 86 | 8 | Pres vif (quaver = 160). Pulsating semiquavers, occasional 'blasts' of acciaccatura quaver. |
| | 2 | 2 | | 60 | 11 | Modéré (quaver = 138). Expressing |
| | 3 | 3 | | 71 | 15 | Un peu plus vif (quaver = 160). Quaver trills in H, RH semiquaver triplet |
| | | 4 | | 71 | 17 | Bien modéré (mais de plus en plus véhément crotchet tied to semiquaver = 58. Repetitive 5 semiquaver, (2+2+1) falling motif, occasionally interrupted by accented slower semiquaver chords |
| | | 5 | | 42 | 10 | Très modéré, Tempo rubato (quaver = 104). Accented 4 note chords in an arching kind of manner |
| | | 6 | | 70 | 10 | Modéré (quaver = 138). Quaver, preceded by 5 semiquaver, and burst scalic upward patterns, followed by semiquaver pattern |
| | | 7 | | 41 | 20 | Similar to section 5. |
| | | 8 | | 81 | 14 | Similar to section 1. |
| | | 9 | | 37 | 29 | Starts similar to Section 5 and 7, followed by trills in LH, hemidemisemiquaver pulsations in RH. |

8
9

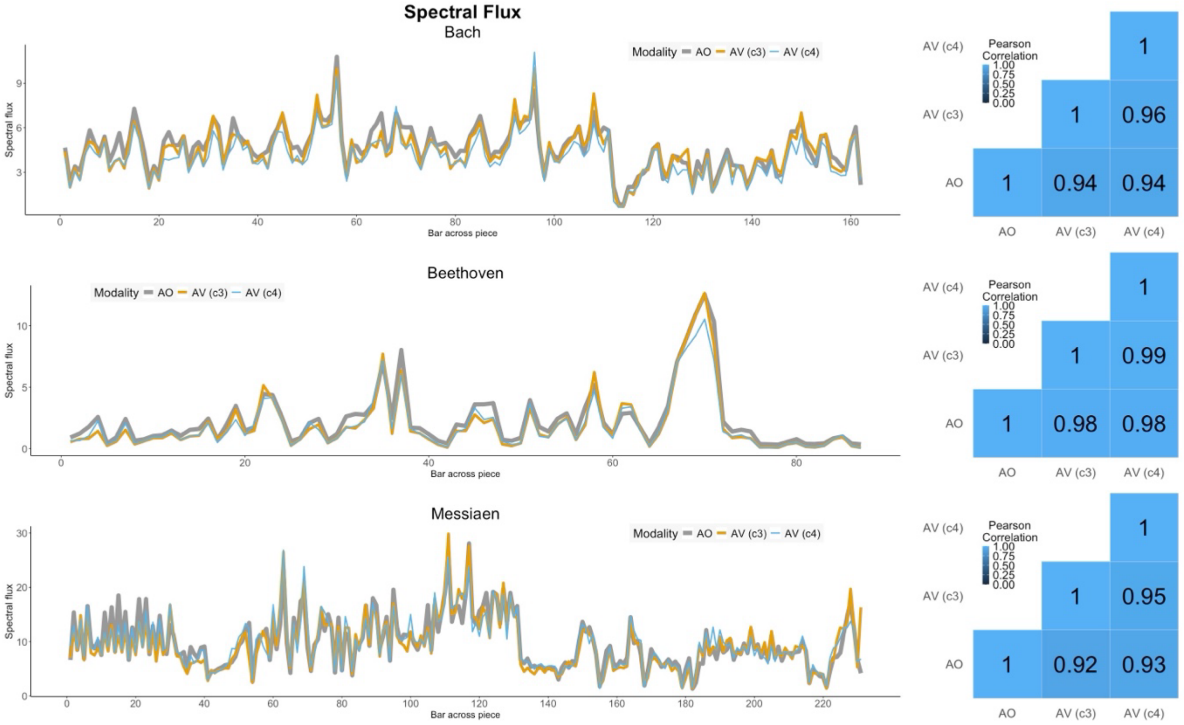

**Supplementary Figure 1.** Assessing similarity of spectral flux across different modality performances for audio-only (AO) performances and audio-visual (AV) performances for concert 3 (c3) and concert 4 (c4). Note: this study was part of an experimental concert series (see Czepiel et al., 2023), where physiology was only collected in concerts 3 and 4. Left panels show the time series and right panels shows correlation matrices of Bach, Beethoven, and Messiaen performances (different rows).

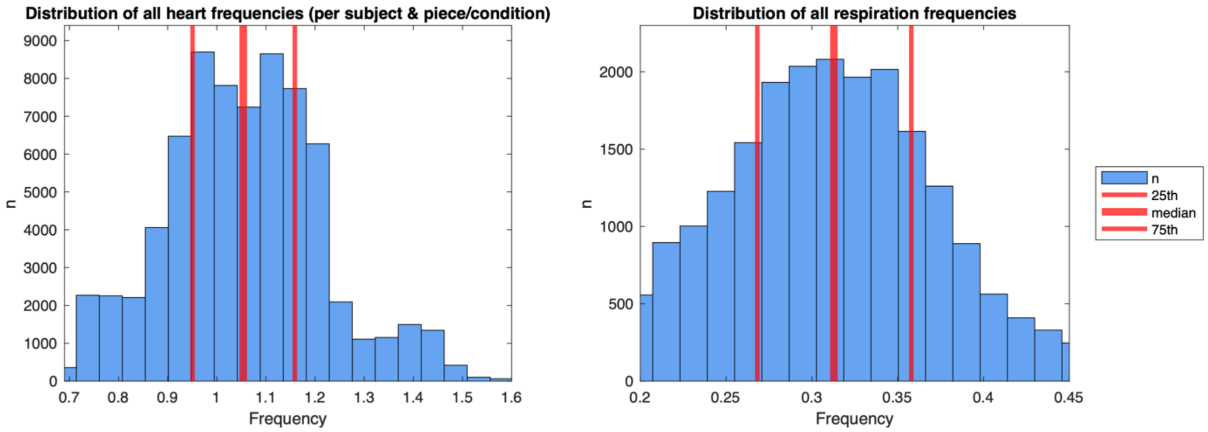

**Supplementary Figure 2.** Distributions of all heart (left panel) and respiration (right panel) frequencies. Frequency is shown on the x-axis, with the number of observations at a certain frequency shown in the blue bars. Red lines show the interquartile range: the thickest red line shows the median frequency, and the outer (thinner) red lines denote the 25th and 75th percentile.

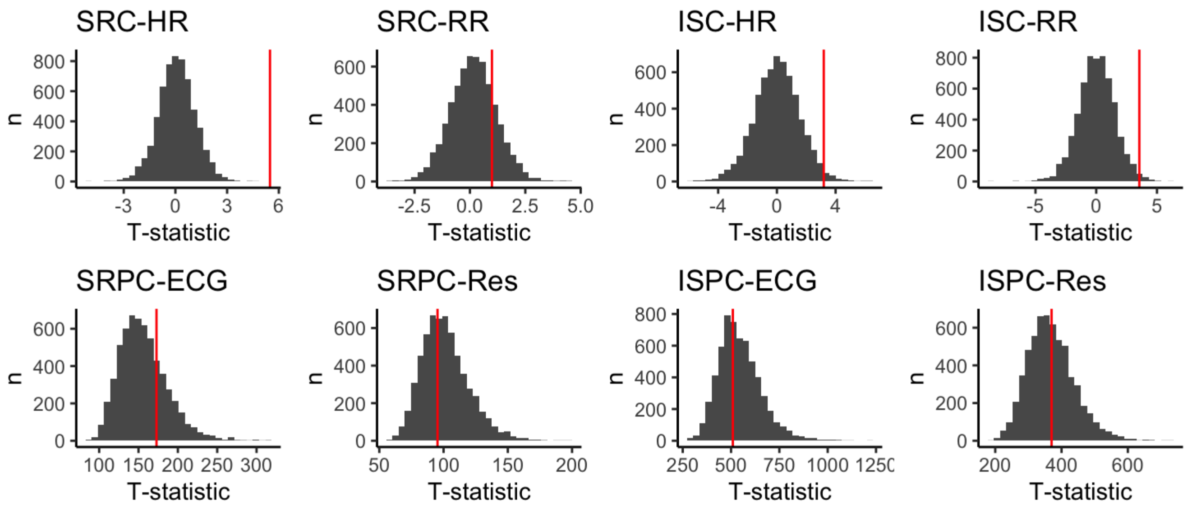

**Supplementary Figure 3.** Distributions of permutation values in grey for all eight of the synchrony values. The red line shows the location of the true test statistic of the time-locked data.

**Supplementary Table 2.** P values from permutation analysis for 30 second sections. (False Discovery Rate, FDR adjusted).

| | Heart | Respiration |
|---|---|---|
| SRC | < .001 | .436 |
| ISC | .065 | .051 |
| SRPC | .436 | .632 |
| ISPC | .632 | .632 |

**Supplementary Table 3.** P values from permutation analysis across different pieces (FDR adjusted).

| | Heart | | | | Respiration | | | |
|---|---|---|---|---|---|---|---|---|
| | SRC-HR | ISC-HR | SRPC-ECG | ISPC-ECG | SRC-RR | ISC-RR | SRPC-Res | ISPC-Res |
| Bach | <.001 | .271 | .239 | .421 | .300 | .224 | .329 | .327 |
| Beethoven | <.001 | .019 | .406 | .697 | .073 | .010 | .805 | .805 |
| Messiaen | <.001 | .016 | .796 | .824 | .803 | .016 | .824 | .580 |

**Supplementary Table 4.** P values from permutation analysis between different modalities (FDR adjusted).

| | Heart | | | | Respiration | | | |
|---|---|---|---|---|---|---|---|---|
| | SRC-HR | ISC-HR | SRPC-ECG | ISPC-ECG | SRC-RR | ISC-RR | SRPC-Res | ISPC-Res |
| Audio-only | <.001 | .145 | .500 | .618 | .394 | .016 | .588 | .534 |
| Audio-visual | <.001 | .023 | .337 | .632 | .337 | .039 | .660 | .632 |

**Supplementary Table 5.** P values from permutation analysis where audience members rated higher than mid-point for 'urge to move' (i.e., a rating of 4 and above on a 7-point scale, see Czepiel et al., 2023).

| | Heart | | | | Respiration | | | |
|---|---|---|---|---|---|---|---|---|
| | SRC-HR | ISC-HR | SRPC-ECG | ISPC-ECG | SRC-RR | ISC-RR | SRPC-Res | ISPC-Res |
| 'Urge to Move' ratings >4 | .016 | .108 | .376 | .376 | .376 | .061 | .376 | .493 |

**Supplementary Table 6.** Linear mixed models comparing synchrony between modalities for maximal model (all random effects specified). These maximal models gave errors (see accompanying code).

| | **SRC-HR** | | | **ISC-HR** | | | **ISC-RR** | | |
|---|---|---|---|---|---|---|---|---|---|
| *Predictors* | *Estimates* | *CI* | *p* | *Estimates* | *CI* | *p* | *Estimates* | *CI* | *p* |
| (Intercept) | 0.050 | 0.020 – 0.080 | **0.001** | 0.012 | 0.002 – 0.022 | **0.020** | 0.021 | 0.009 – 0.033 | **0.001** |
| mod [AV] | 0.009 | -0.006 – 0.024 | 0.246 | 0.010 | 0.001 – 0.019 | **0.028** | 0.002 | -0.009 – 0.013 | 0.698 |
| **Random Effects** | | | | | | | | | |
| $\sigma^2$ | 0.01 | | | 0.00 | | | 0.01 | | |
| $\tau_{00}$ | $0.00_{\text{mac:conc}}$ | | | $0.00_{\text{mac:conc}}$ | | | $0.00_{\text{mac:conc}}$ | | |
| | $0.00_{\text{piece}}$ | | | $0.00_{\text{piece}}$ | | | $0.00_{\text{piece}}$ | | |
| | $0.00_{\text{conc}}$ | | | $0.00_{\text{conc}}$ | | | $0.00_{\text{conc}}$ | | |
| $\tau_{11}$ | $0.00_{\text{mac1.modAO}}$ | | | $0.00_{\text{mac1.modAO}}$ | | | $0.00_{\text{mac1.modAO}}$ | | |
| | $0.00_{\text{mac2.modAV}}$ | | | $0.00_{\text{mac2.modAV}}$ | | | $0.00_{\text{mac2.modAV}}$ | | |
| $\rho_{01}$ | | | | | | | | | |
| $\rho_{01}$ | | | | | | | | | |
| N | $2_{\text{conc}}$ | | | $2_{\text{conc}}$ | | | $2_{\text{conc}}$ | | |
| | $3_{\text{piece}}$ | | | $3_{\text{piece}}$ | | | $3_{\text{piece}}$ | | |
| | $16_{\text{mac}}$ | | | $16_{\text{mac}}$ | | | $16_{\text{mac}}$ | | |
| Observations | 1075 | | | 1075 | | | 1152 | | |
| Marginal $R^2$ / Conditional $R^2$ | 0.002 / NA | | | 0.008 / NA | | | 0.000 / NA | | |

**Supplementary Table 7.** Linear mixed models for modality and time window for temporal epochs centred around section boundaries for maximal model (all random effects specified). These maximal models gave errors (see accompanying code).

| | SRC-HR | | | ISC-HR | | | ISC-RR | | |
|---|---|---|---|---|---|---|---|---|---|
| *Predictors* | *Estimates* | *CI* | *p* | *Estimates* | *CI* | *p* | *Estimates* | *CI* | *p* |
| (Intercept) | 0.04 | 0.02 – 0.06 | **<0.001** | 0.02 | 0.01 – 0.03 | **<0.001** | 0.05 | 0.03 – 0.07 | **<0.001** |
| mod [AV] | 0.02 | 0.00 – 0.04 | **0.017** | 0.02 | 0.01 – 0.03 | **0.001** | -0.03 | -0.06 – -0.00 | **0.038** |
| win-10 | -0.01 | -0.03 – 0.01 | 0.339 | -0.01 | -0.02 – -0.00 | **0.014** | -0.00 | -0.03 – 0.03 | 0.828 |
| win-5 | 0.01 | -0.01 – 0.03 | 0.356 | -0.00 | -0.01 – 0.01 | 0.635 | -0.00 | -0.03 – 0.02 | 0.792 |
| win [5] | -0.03 | -0.05 – -0.00 | **0.018** | -0.01 | -0.02 – 0.00 | 0.059 | -0.04 | -0.06 – -0.01 | **0.008** |
| win [10] | -0.03 | -0.05 – -0.00 | **0.017** | -0.01 | -0.02 – -0.00 | **0.013** | -0.03 | -0.06 – -0.01 | **0.016** |
| modAV:win-10 | -0.00 | -0.03 – 0.02 | 0.800 | -0.01 | -0.02 – 0.01 | 0.226 | 0.03 | -0.01 – 0.07 | 0.155 |
| modAV:win-5 | 0.00 | -0.02 – 0.03 | 0.714 | -0.00 | -0.02 – 0.01 | 0.602 | 0.01 | -0.02 – 0.05 | 0.490 |
| mod [AV] × win [5] | -0.02 | -0.05 – 0.00 | 0.067 | -0.02 | -0.03 – -0.01 | **0.002** | 0.04 | 0.00 – 0.08 | **0.031** |
| mod [AV] × win [10] | -0.03 | -0.06 – -0.01 | **0.005** | -0.02 | -0.04 – -0.01 | **0.001** | 0.04 | -0.00 – 0.07 | 0.056 |
| **Random Effects** | | | | | | | | | |
| $\sigma^2$ | 0.00 | | | 0.00 | | | 0.01 | | |
| $\tau_{00}$ | $0.00_{\text{mac.conc}}$ | | | $0.00_{\text{mac.conc}}$ | | | $0.00_{\text{mac.conc}}$ | | |
| | $0.00_{\text{piece}}$ | | | $0.00_{\text{piece}}$ | | | $0.00_{\text{piece}}$ | | |
| | $0.00_{\text{conc}}$ | | | $0.00_{\text{conc}}$ | | | $0.00_{\text{conc}}$ | | |
| $\tau_{11}$ | $0.00_{\text{mac.win10}}$ | | | $0.00_{\text{mac.win-10}}$ | | | $0.00_{\text{mac.win-10}}$ | | |
| | $0.00_{\text{mac.1.modAV}}$ | | | $0.00_{\text{mac.win-5}}$ | | | $0.00_{\text{mac.win-5}}$ | | |
| | $0.00_{\text{mac1.win0}}$ | | | $0.00_{\text{mac.win5}}$ | | | $0.00_{\text{mac.win5}}$ | | |
| | $0.00_{\text{mac2.win-10}}$ | | | $0.00_{\text{mac.win10}}$ | | | $0.00_{\text{mac.win10}}$ | | |
| | $0.00_{\text{mac3.win-5}}$ | | | $0.00_{\text{mac.1.modAV}}$ | | | $0.00_{\text{mac1.win0}}$ | | |
| | $0.00_{\text{mac4.win5}}$ | | | $0.00_{\text{mac1.win0}}$ | | | $0.00_{\text{mac2.win-10}}$ | | |
| | $0.00_{\text{mac5.win10}}$ | | | $0.00_{\text{mac2.win-10}}$ | | | $0.00_{\text{mac3.win-5}}$ | | |
| | $0.00_{\text{mac.modAO}}$ | | | $0.00_{\text{mac3.win-5}}$ | | | $0.00_{\text{mac4.win5}}$ | | |
| | $0.00_{\text{mac.modAV}}$ | | | $0.00_{\text{mac4.win5}}$ | | | $0.00_{\text{mac5.win10}}$ | | |
| | | | | $0.00_{\text{mac5.win10}}$ | | | $0.00_{\text{mac.modAO}}$ | | |
| | | | | $0.00_{\text{mac.modAO}}$ | | | $0.00_{\text{mac.modAV}}$ | | |
| | | | | $0.00_{\text{mac.modAV}}$ | | | | | |
| $\varrho_{01}$ | | | | | | | | | |
| $\varrho_{01}$ | | | | | | | | | |
| N | $2_{\text{conc}}$ | | | $2_{\text{conc}}$ | | | $2_{\text{conc}}$ | | |
| | $3_{\text{piece}}$ | | | $3_{\text{piece}}$ | | | $3_{\text{piece}}$ | | |
| | $16_{\text{mac}}$ | | | $16_{\text{mac}}$ | | | $16_{\text{mac}}$ | | |
| Observations | 645 | | | 645 | | | 690 | | |
| Marginal $R^2$ / Conditional $R^2$ | 0.171 / NA | | | 0.133 / NA | | | 0.027 / NA | | |

**Supplementary Table 8.** Linear mixed models of heart and respiration rate (HR, RR), at time window for temporal epochs centred around section boundaries for maximal model (all random effects specified). These maximal models gave errors (see accompanying code).

| | Heart Rate | | | Respiration Rate | | |
|---|---|---|---|---|---|---|
| *Predictors* | *Estimates* | *CI* | *p* | *Estimates* | *CI* | *p* |
| (Intercept) | 61.79 | 52.18 – 71.40 | **<0.001** | 18.80 | 17.49 – 20.12 | **<0.001** |
| win-10 | 0.45 | 0.06 – 0.85 | **0.024** | 0.15 | -0.13 – 0.44 | 0.285 |
| win-5 | 0.25 | -0.14 – 0.64 | 0.213 | -0.24 | -0.52 – 0.05 | 0.104 |
| win [5] | 0.37 | -0.02 – 0.76 | 0.064 | 0.10 | -0.18 – 0.39 | 0.474 |
| win [10] | 0.53 | 0.14 – 0.92 | **0.008** | 0.07 | -0.21 – 0.35 | 0.628 |
| **Random Effects** | | | | | | |
| $\sigma^2$ | 2.54 | | | 1.45 | | |
| $\tau_{00}$ | 45.31 mac.conc | | | 6.65 mac.conc | | |
| | 0.15 piece | | | 0.44 piece | | |
| | 42.36 conc | | | 0.00 conc | | |
| $\tau_{11}$ | 9.47 mac1.win0 | | | 0.00 mac.win-10 | | |
| | 10.05 mac2.win-10 | | | 0.00 mac.win-5 | | |
| | 9.81 mac3.win-5 | | | 0.00 mac.win5 | | |
| | 9.86 mac4.win5 | | | 0.00 mac.win10 | | |
| | 9.40 mac5.win10 | | | 0.00 mac1.win0 | | |
| | 6.79 mac.modAO | | | 0.00 mac2.win-10 | | |
| | 16.86 mac.modAV | | | 0.00 mac3.win-5 | | |
| | | | | 0.00 mac4.win5 | | |
| | | | | 0.00 mac5.win10 | | |
| | | | | 1.49 mac.modAO | | |
| | | | | 0.38 mac.modAV | | |
| $\rho_{01}$ | | | | | | |
| $\rho_{01}$ | | | | | | |
| N | 2 conc | | | 2 conc | | |
| | 3 piece | | | 3 piece | | |
| | 16 mac | | | 16 mac | | |
| Observations | 645 | | | 690 | | |
| Marginal $R^2$ / Conditional $R^2$ | 0.013 / NA | | | 0.013 / NA | | |
